# Replication stress response in fission yeast differentially depends on maintaining proper levels of Srs2 helicase and Rrp1, Rrp2 DNA translocases

**DOI:** 10.1101/2023.12.05.570082

**Authors:** Gabriela Baranowska, Dorota Misiorna, Wojciech Białek, Karol Kramarz, Dorota Dziadkowiec

**Author notes:** Corresponding authors: (DD), (KK).

## Abstract

Homologous recombination is a key process that governs the stability of eukaryotic genomes during DNA replication and repair. Multiple auxiliary factors regulate the choice of homologous recombination pathway in response to different types of replication stress. Using *Schizosaccharomyces pombe* we have previously suggested the role of DNA translocases Rrp1 and Rrp2, together with Srs2 helicase, in the common synthesis dependent strand annealing sub-pathway of homologous recombination. Here we show that all three proteins are important for completion of replication after hydroxyurea exposure and provide data comparing the effect of overproduction of Srs2 with Rrp1 and Rrp2. Upregulation of Srs2 protein levels leads to enhanced replication stress, chromosome instability and viability loss, as previously reported for Rrp1 and Rrp2. Interestingly, our data suggests that dysregulation of Srs2, Rrp1 and Rrp2 protein levels differentially affects checkpoint response. Overproduction of Srs2 activates simultaneously DNA damage and replication stress response checkpoints, while cells overproducing Rrp1 mainly launch DNA damage checkpoint. Upregulation of Rrp2 primarily leads to replication stress response checkpoint activation. Overall, we propose that Srs2, Rrp1 and Rrp2 have important and independent functions for maintenance of distinct difficult to replicate regions of the genome.

## Introduction

Homologous recombination (HR) is crucial for maintaining genome stability due to its involvement in the repair of DNA double-strand breaks (DSBs) and recovery of arrested replication forks. In *S. pombe* the key Rad51 recombinase is assisted by auxiliary factors that regulate its activity, and are evolutionary conserved not only in *S. cerevisiae* but also in humans [1]. Among them two complexes, Rad55-Rad57 and Swi5-Sfr1, were proposed to act in parallel to stabilize the Rad51 nucleofilament and stimulate Rad51-mediated strand exchange [2]. The choice of specific complex may determine the pathway and thus the final outcomes of HR, since it has been shown that second-end capture and crossover (CO) production in *S. pombe* is dependent on Rad55-Rad57, but does not require Swi5-Sfr1 [3]. We have previously described in *S. pombe* another complex, Rrp1-Rrp2 [4], and proposed it to function together with Swi5-Sfr1 and Srs2 helicase, but independently of Rad55-Rad57, in a synthesis dependent strand annealing pathway (SDSA) of HR [5]. Thus, according to our model, in the absence of Rad57-dependent branch of HR, Rad51 generated recombination products would be channelled into SDSA, away from the pathway requiring dissolution by Rqh1. Indeed, we have observed that the presence of Rrp1 and Rrp2 was required for the full rescue of the *rqh1*Δ mutant’s HU sensitivity and aberrant mitosis phenotype by deletion of *rad57+* [5].

Studies of the effect of upregulation of Rrp1 and Rrp2 protein levels allowed to identify their novel biological functions in genome stability maintenance [6]. We have shown that Rrp1, and to a lesser degree Rrp2, are involved in the modulation of histone levels and their overproduction resulted in defects in centromere structure and function [7]. Based on the discovery that Rrp2 can protect SUMOylated Top2 from premature degradation [8], we have proposed that Rrp2 may protect telomeres against Top2-induced DNA damage, a function it does not share with Rrp1 [7]. Recently, we have demonstrated that Rrp1, but not Rrp2, can protect the cells from the toxicity of *rad51*+ overexpression [9]. In addition, our *in vitro* and *in vivo* data showed that Rrp1 is a DNA-dependent translocase and ubiquitin ligase that can regulate Rad51 binding to dsDNA [9].

In summary, even though Rrp1 and Rrp2 have been established to act as a complex in replication stress response within a sub-pathway of HR, they can also have functions that are, at least partially, independent from each other.

Srs2 is a DNA-dependent helicase that belongs to the UvrD-like superfamily present in S. *cerevisiae* and *S. pombe*, with no direct homologue in human cells. Most of the information on Srs2 important roles in DNA replication, recombination and repair comes from work in *S. cerevisiae*, where it has been very extensively studied [10]. Several helicases, such as WRN, HELB, or PARI are regarded as potential functional human Srs2 orthologues, fulfilling some of the functions attributed to Srs2 in *S. cerevisiae*. Srs2 has been first described as “anti-recombinase” due to its ability to remove Rad51 from ssDNA [11,12]. Nevertheless, Srs2 has also been demonstrated to remove single strand DNA binding protein, RPA, from DNA and thus regulate checkpoint response [13,14]. Srs2 has also been shown to inhibit homologous recombination during post-replication repair in *S. cerevisiae* through its interaction with PCNA [15], although this activity seems not to be conserved in *S. pombe* [16]. Later however it has been proposed that Srs2 is rather involved in the determination of repair pathway choice and directs the processing of HR intermediates into the SDSA by several mechanisms unrelated to its interaction with Rad51: inhibiting D-loop extension through competition with delta polymerase, as well as direct D-loop unwinding [17,18]. Srs2 clearly is a multifunctional protein in *S. cerevisiae* and still more roles will probably be ascribed to it in the future. It is important to establish, which of these functions are conserved in *S. pombe*, and which have been transferred to other proteins exhibiting helicase/translocase activities. This clarification would considerably help to elucidate the division of labour between functional orthologues of Srs2 in humans.

## Materials and methods

### Yeast strains, plasmids and general methods

Strains, plasmids and primers used in this study are listed in Tables S1, S2 and S3, respectively. Media used for *S. pombe* growth were as described [27]. Yeast cells were grown in complete yeast extract plus supplements (YES) medium or glutamate supplemented Edinburgh minimal medium (EMM) at 28°C. Thiamine at 5 μg/mL, geneticin (ICN Biomedicals) at 100 μg/mL and nurseotricin (Werner Bioagents) at 200 μg/mL were added where required. Multiple mutants were obtained by genetic crossing of relevant single mutants followed either by random spore analysis or by tetrad dissection. pREP81-FLAG plasmid carrying wild-type *srs2*^+^ was constructed using the Gibson Assembly® Cloning Kit (NEB E5510S). After Gibson cloning, the construct was confirmed by sequencing and NdeI and SmaI digested insert was cloned into other required plasmids.

### Spot assays

Cells were grown to mid-log phase, then serially diluted by 10-fold and 2 µL aliquots were spotted onto relevant plates (YES or EMM) without drugs or plates containing camptothecin (CPT) or hydroxyurea (HU). Plates were incubated for 3-5 days at 28°C and photographed. All assays were repeated at least twice.

### Survival assays

Cells were grown for 48 hours in minimal medium with (repressed conditions) or without thiamine (overexpression) at 28°C, serially diluted, plated onto YES medium and incubated for 3-5 days at 28°C. The viable cells were counted and the percentage of survival for gene overexpression conditions was calculated against the repressed control.

### Yeast two-hybrid assay

Gal4-based Matchmaker Two-Hybrid System 3 (Clontech) was used according to the manufacturer’s instructions. The indicated proteins were fused to the GAL4 activation domain (AD) in pGADT7 vector and the GAL4 DNA-binding domain (DBD) in pGBKT7, and expressed in the *S. cerevisiae* strain AH109. Transformants were selected on synthetic dextrose drop-out medium without Leu and Trp (SD DO-2), and then spotted on SD DO-2 as control and high stringency medium without Leu, Trp, His and Ade (SD DO-4). Plates were incubated for 3-5 days at 28°C and photographed.

### Fluorescence microscopy

To determine the localization of overproduced EGFP-Srs2 and the influence of *srs2*^*+*^ overexpression on the accumulation of Rad11-EGFP foci, appropriate transformants were grown for 24 h in EMM medium without thiamine. 1 mL of culture was harvested, washed with water and subjected to fluorescent microscopy analysis. For examination of mitotic defects induced by *srs2*^+^ overexpression, samples taken from respective transformant cultures grown for 48 hours in EMM medium without thiamine were fixed in 70% ethanol. After rehydration, cells were stained with 1 mg/mL 4’,6-diamidino-2-phenylindole (DAPI) and 1 mg/mL p-phenylenediamine in 50% glycerol and examined by fluorescence microscopy. Images were captured under 100x magnification using Axio Imager A.2 (Carl Zeiss) and analysed with Axiovision rel. 4.8.

### Chromosome loss

Single white colonies from indicated transformants grown on EMM low Ade plates (adenine concentration 7.5 mg/L) with thiamine were inoculated into EMM medium without thiamine and incubated for 48 h at 28°C. Then cultures were diluted, plated on YES low Ade plates and incubated for 3-4 days at 28°C. The percentage of red to white colonies was then calculated as a readout for the loss of unessential mini-chromosome.

### Pulse field gel electrophoresis

Logarithmic yeast cultures (grown in rich YES medium) were diluted to OD_600_ ∼0.5, then exposed to 20 mM HU for 4 hours, subsequently washed with water and released to fresh YES medium. At indicated time points 20 mL of cell culture was collected, washed with cold 50 mM EDTA pH 8 and digested with litycase (Sigma, L4025) in CSE buffer (20 mM citrate/phosphate pH 5.6, 1.2 M sorbitol, 40 mM EDTA pH 8). Spheroplasts were embedded into 1% UltraPure™ Agarose (Invitrogen, 16500) and put into 4 agarose plugs per each time point. Obtained plugs were incubated in Lysis Buffer 1 (50 mM Tris-HCl pH 7.5, 250 mM EDTA pH 8, 1 % SDS) for 90 minutes in 55°C and then digested in Lysis Buffer 2 (1 % N-lauryl sarcosine, 0.5 M EDTA pH 9.5, 0.5 mg/mL proteinase K) o/n at 55°C. Lysis Buffer 2 was changed next morning for fresh one and digestion was continued o/n at 55°C. Plugs were then stored at 4°C. Pulse field gel electrophoresis was carried out on Biorad CHEF-DR-III system for 48 hours at 2.0 V/cm, with an angle 120°, at 14°C. Single switch time was set at 1800 s, pump speed 70. Electrophoresis was carried out in 1x TAE buffer. Chromosomes were visualized at Biorad Chemidoc MP after gel staining in ethidium bromide (10 μg/mL) for 30 min and washing for 30 min in 1x TAE.

### Whole protein extract analysis

The trichloroacetic acid (TCA) method was used to obtain protein extracts. Mid-logarithmic cultures (∼10^8^) of indicated strains were harvested after 24 hour of induction of the *nmt* promoter by removal of thiamine from media and lysed with lysis buffer (2 M NaOH, 7% β-mercaptoethanol). Total protein was precipitated with 50% TCA. Pellet was resuspended in 1 M Tris at pH 8 and 4x Laemmli buffer was added (250 mM Tris-HCl, pH 6.8, 8% SDS, 20% glycerol, 0.02% Bromophenol blue, 7% β-mercaptoethanol). Samples were analysed by SDS-PAGE and Western blotting using anti-FLAG (Sigma-Aldrich, F1804), anti-H2A (Active Motif, 39235), anti-γ-H2A (Abcam, ab15083), anti-HA (Roche, 11538816001) antibodies. Antibodies or Ponceau S staining (Sigma-Aldrich) of blotted membranes were used as loading controls. Image Lab (Western blots) or ImageJ software (Ponceau S staining) were used for protein quantification. Relative intensity was calculated by dividing sample intensities by the mean of control intensities obtained for each blot (details for each experiment are provided in figure captions). For each experiment data from at least two different transformants from two independent protein isolations were analysed.

### Checkpoint activation

Cells were grown for 48 hours in EMM minimal medium without thiamine (over-expression conditions) at 28°C. 1 mL samples were taken from each culture and subjected to microscopy analysis in order to determine cell length. Images were captured under 100x magnification using Axio Imager A.2 (Carl Zeiss) and analysed with Axiovision rel. 4.8. From the remaining cultures whole cell protein extracts were isolated and subjected to Western blot analysis described above in order to determine levels of H2A and Chk1 phosphorylation.

### Statistical data analysis

Student’s t test was used to calculate the P-values (**** *p* ≤ 0.0001, *** *p* ≤ 0.001, ** 0.001 < *p* ≤ 0.01, * 0.01 < *p* ≤ 0.05).

## Results

### Srs2 helicase may have a function in a similar HR repair pathway to Rrp1 and Rrp2

We have previously demonstrated that Rrp1 and Rrp2 are involved in replication stress response and may act together with Srs2 helicase within Swi5-Sfr1 branch of synthesis-dependent strand annealing homologous recombination repair pathway [5]. Additionally we found that the presence of Rrp1 and Rrp2 was required for the full rescue of the *rqh1*Δ mutant’s HU sensitivity and aberrant mitosis phenotype by deletion of *rad57+* [5]. Here we show that the deletion of *srs2*+ also increases the double *rad57*Δ*rqh1*Δ mutant’s HU sensitivity, to the same level as the deletion of *rrp1+* or *rrp2+* (Fig.1A). Furthermore, the incidence of aberrant mitotic phenotypes, such as cut or non-disjunction nuclei, is increased in *rad57*Δ*rqh1*Δ*srs2*Δ mutant compared to *rad57*Δ*rqh1*Δ (Fig.1B), similar to what has been shown earlier for *rad57*Δ*rqh1*Δ*rrp1*Δ and *rad57*Δ*rqh1*Δ*rrp2*Δ [5]. This lends support to our published model placing Rrp1, Rrp2 and Srs2 in a common pathway.

**Fig 1.**
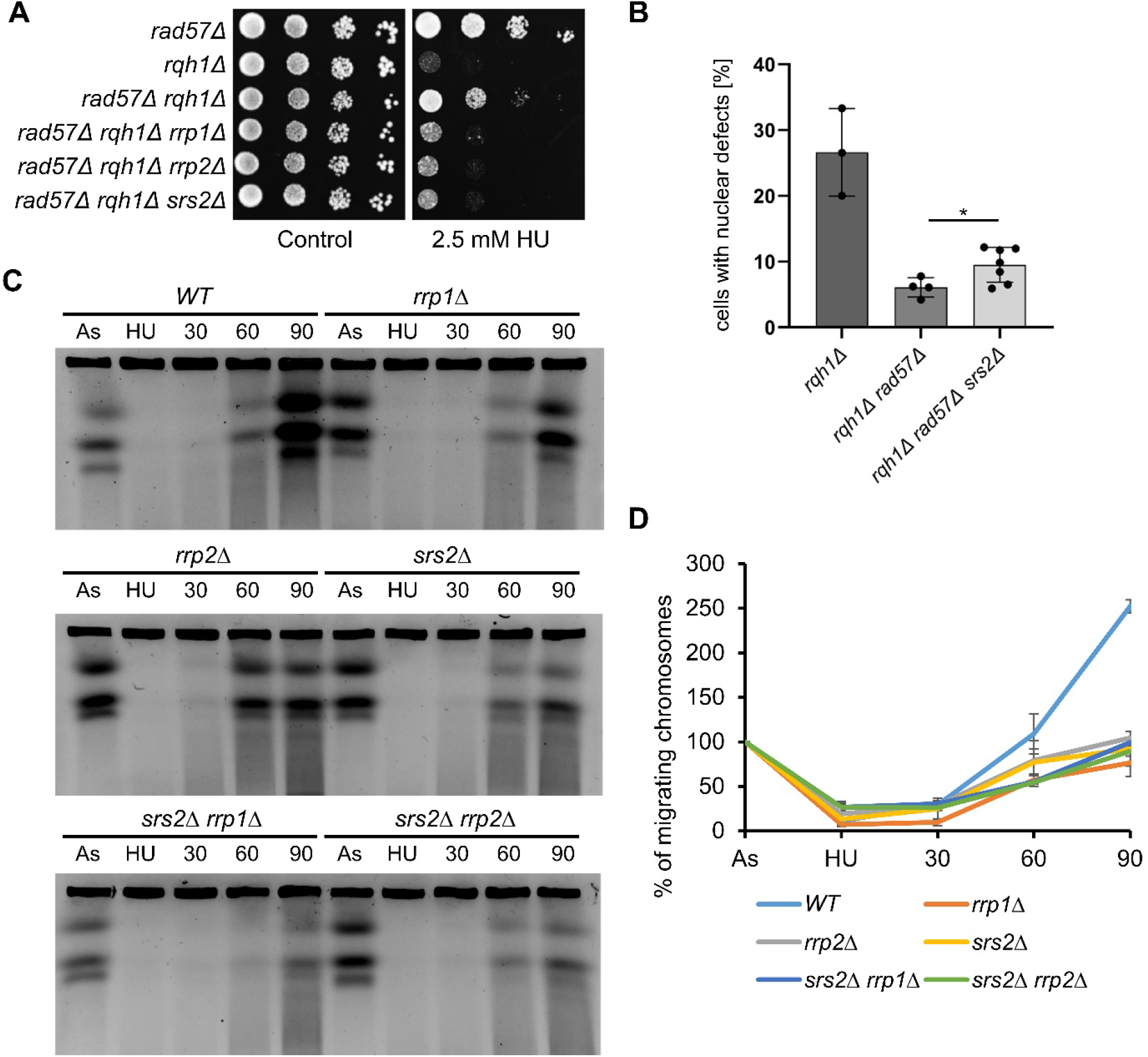
Srs2 helicase shares some functions in HR with Rrp1 and Rrp2 translocases. Srs2 helicase is required for the rescue of *rqh1*Δ HU sensitivity (A) and aberrant mitosis (B) by deletion of *rad57*+. Cells were appropriately diluted and spotted on YES plates with HU, incubated for 4 days and photographed. Mitotic aberrations were observed by DAPI staining of the nuclei of respective mutants grown for 24 h in complete medium. At least three independent cultures of each strain were examined. Error bars represent the standard deviation about the mean values. Student’s t-test was performed to calculate P-values (* 0.01 < P ≤ 0.05). (C) Chromosome analysis by pulse field gel electrophoresis (PFGE) in respective mutants in indicated time points: As – asynchronous, logarithmic cells exposed to 20 mM HU by 4 hours (HU) and after drug removal released into fresh rich medium YES at 30oC to verify ability to resolve replication intermediates (30, 60, 90 timepoints). (D) A graph presenting quantification of % of chromosomes migrating in gel from indicated timepoints. Values are means of two independent biological experiments. Error bars are standard deviation (SD).

We have previously shown that while cells devoid of *srs2+* were slightly sensitive to hydroxyurea (HU) the single mutants *rrp1*Δ and *rrp2*Δ were not [5]. However, when the ability to resume replication upon transient treatment of studied mutants with 20 mM HU was examined by pulse field gel electrophoresis we found that not only the presence of Srs2 but also of Rrp1 and Rrp2 was required for proper replication completion (Fig. 1C). While wild-type (WT) cells completed replication 90 minutes after incubation in drug-free medium, indicated by the doubling of the intensity of chromosomes compared to initial asynchronous culture, in both *rrp1*Δ and *rrp2*Δ mutants the increase in chromosomes intensities at 90 min time point was far less pronounced (Fig. 1C). The double mutants simultaneously devoid of *srs2+* and *rrp1+* or *rrp2+* did not show any further delay in replication completion, suggesting the role of Srs2 in resuming replication after HU block may be independent of Rrp1 and Rrp2 (Fig. 1D).

Previously we have found that even though *rrp1*Δ and *rrp2*Δ single mutants displayed very subtle phenotypes, over-expression of *rrp1+* and *rrp2+* genes allowed to identify novel biological functions for both proteins, at least partially independent from each other [7]. We thus decided to perform similar analysis for Srs2 protein.

### Upregulation of Srs2 protein levels leads to growth defects independent of Rad51, Rrp1 and Rrp2

It has been demonstrated in *S. cerevisiae* that an increase in Srs2 copy number is toxic in certain mutant contexts [19] and we have shown that overproduction of Rrp1 or Rrp2 causes viability loss in otherwise wild-type cells grown under unperturbed conditions [7]. In order to investigate the effect of *srs2*^+^ over-expression on cell growth, we have cloned the *srs2*+ gene under the *nmt* promoter into several pREP expression plasmids (Fig. 2A). The Srs2 protein was produced even when the *nmt* promoter was repressed by addition of thiamine (Fig. 2B) and obtained constructs were able to complement the CPT and HU sensitivity of *srs2*Δ mutant (Fig. 2C). Overproduced EGFP-Srs2 localized to the nucleus and seemed to be enriched in the nucleolus, that contains rDNA repeats (Fig. 2D), consistent with Srs2 being previously shown to be important for rDNA stability [20].

**Fig 2.**
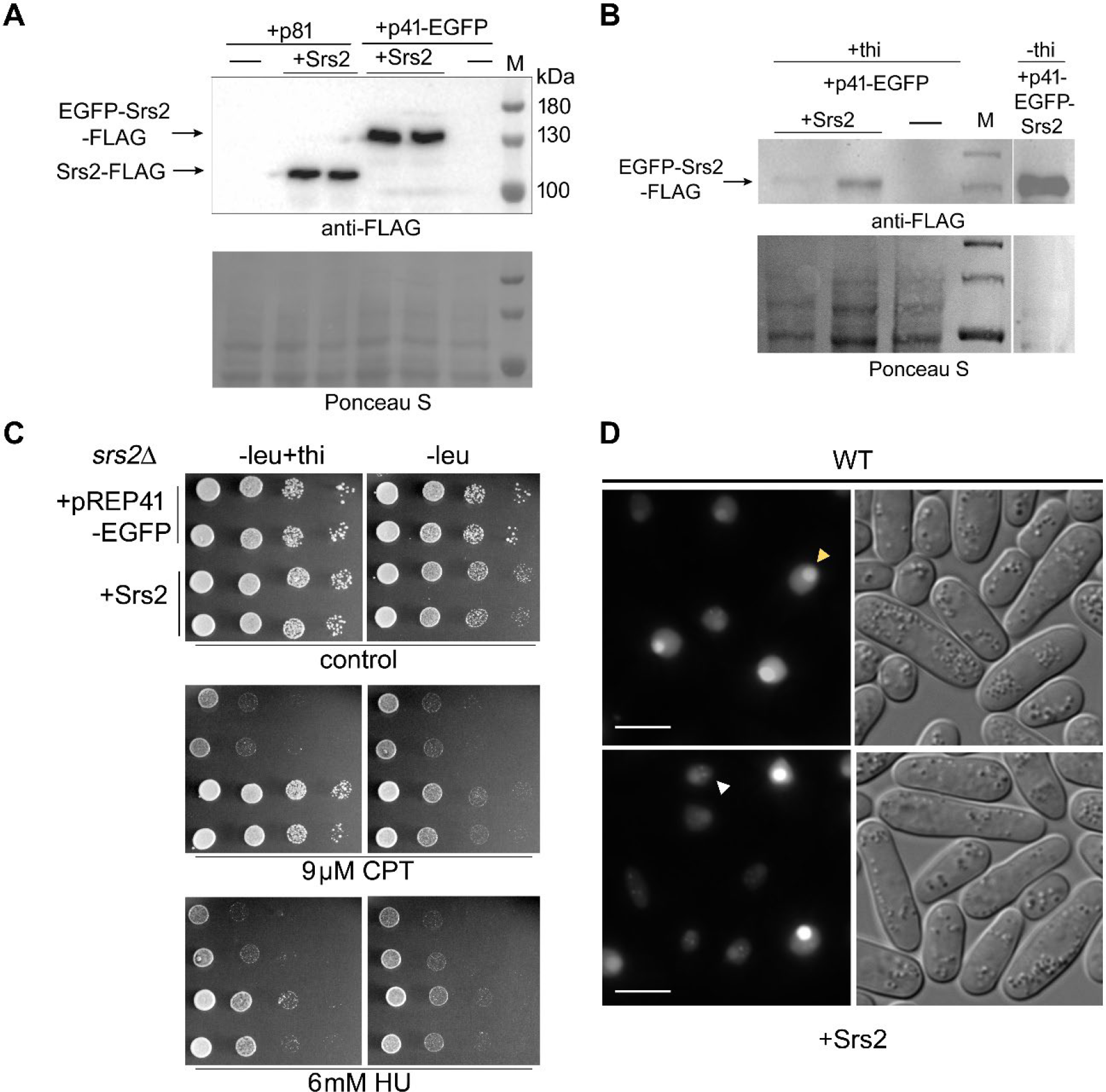
Cloning of *srs2*+ gene. (A) C-terminal FLAG tagged *srs2*+ gene was cloned into several expression plasmids under medium (p41 or 42) or low (p81) strength *nmt* promoter. Western blot analysis with anti-FLAG antibody of protein extracts from transformants containing *srs2-FLAG* plasmids grown in minimal media without thiamine (under expressing inducing conditions, -thi). (B) Residual Srs2-FLAG protein is present even in cultures containing thiamine (+thi) when *nmt* promoter was repressed. (C) Transformation of *srs2*Δ strain with *EGFP*-*srs2-FLAG* expressing plasmid reversed its HU and CPT sensitivity. Cells were appropriately diluted and spotted on EMM plates with HU or CPT supplemented or not with thiamine, incubated for 6 days and photographed. (D) Overproduced EGFP-Srs2-FLAG localises mainly to the nucleolus (yellow arrowhead) and forms spontaneous foci (white arrowhead) in wild type strain. Cells from induced cultures used in (B) were analysed by fluorescence microscopy. Scale bar indicates 10 µm.

The complementation of genotoxin sensitivity was greater on media containing thiamine, when the *nmt* promoter was repressed, resulting in much lower amount of Srs2 produced (Fig. 2B and C, + thi lanes) suggesting that upregulation of Srs2 protein levels might also be toxic in *S. pombe*. Indeed, *srs2*+ over-expression from the medium strength *nmt41* promoter resulted in the growth defect and viability loss, although not as pronounced as those for *rrp2*+, but comparable to those seen for *rrp1*+ (Fig.3A, B) [7]. The toxicity of Srs2 overproduction was not dependent on the presence of Rrp1 and seemed only slightly more pronounced in cells lacking Rrp2 and Rad51 recombinase (Fig. 3C, D). In *S. cerevisiae* it has also been shown that toxicity of Srs2 overproduction was not mediated by Rad51 [19]. Similarly, growth defect induced by *rrp1*+ or *rrp2*+ over-expression in the *srs2*Δ mutant was the same as in wild-type (Fig. 3E). Moreover, our yeast-two-hybrid analysis suggests that Srs2 does not form a direct complex with Rrp1 or Rrp2 (Fig. 3F). The above data indicate that the effects of overexpression of these three genes are independent of each other.

**Fig 3.**
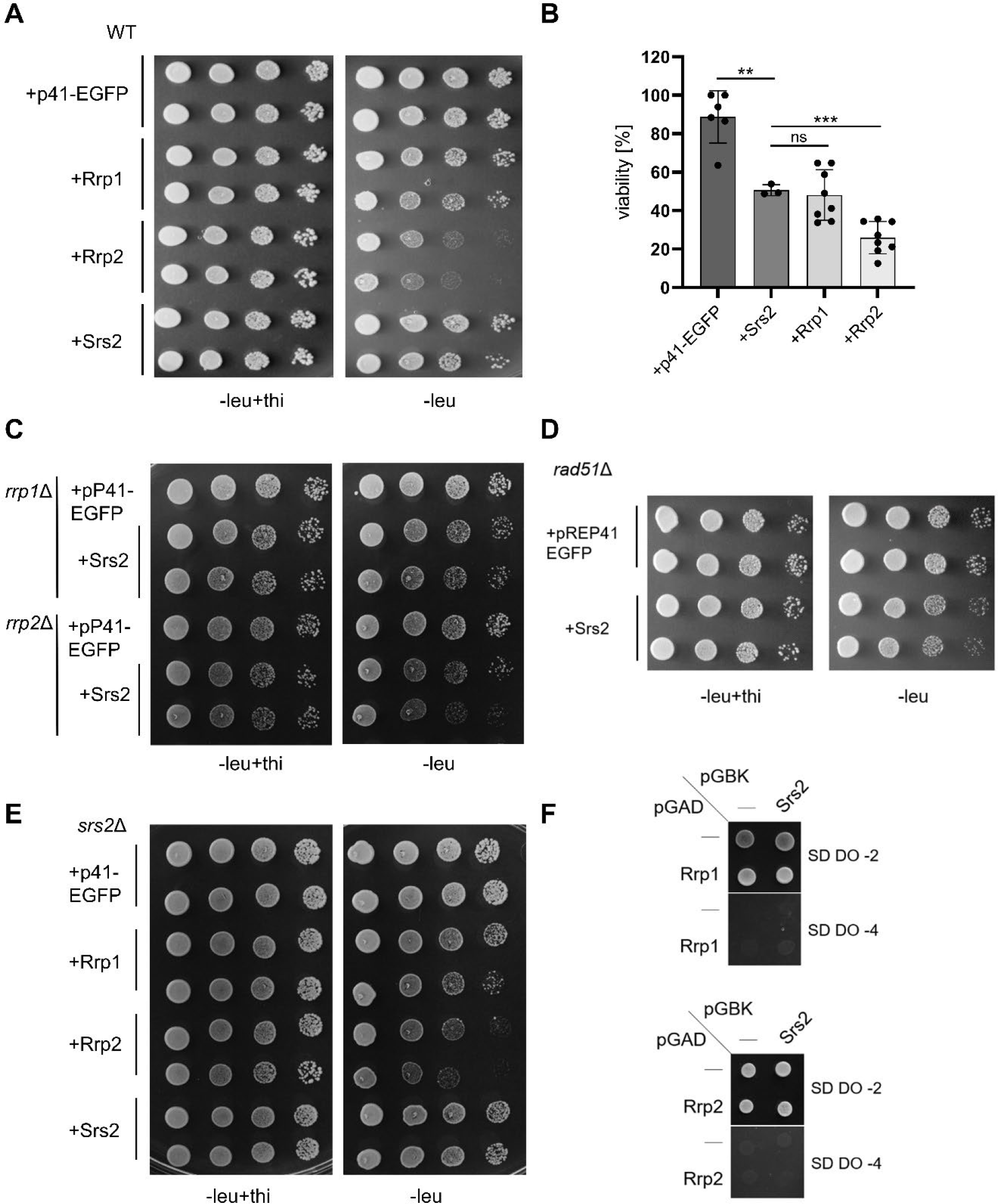
Upregulation of Srs2 protein levels leads to growth defect. Induction of *srs2*+ expression leads to growth defect (A) and viability loss (B) comparable to that seen for *rrp1*+ overexpression. Cells of respective transformants were appropriately diluted and spotted on EMM plates where *srs2*+ expression was induced (-leu) or repressed (-leu + thi), incubated for 6 days and photographed. For viability the ratio of surviving cells of indicated transformants grown under inducing conditions to those grown without induction was determined. The experiment was repeated for at least three independent transformants. Error bars represent the standard deviation about the mean values. Student’s t-test was performed to calculate P-values (*** P ≤ 0.001, ** 0.001 < P ≤ 0.01, ns 0.05≤P). Deletion of *rrp1*+ or*rrp2*+ (C) nor *rad51*+ (D) did not have a major effect on growth defect induced by overproduction of Srs2. (E) Overexpression of *rrp1+, rrp2+* and *srs2+* was not affected by the lack of *srs2+* gene. (F) Srs2 does not interact directly with Rrr1 or Rrp2 as seen in yeast-two hybrid system. Transformants were selected on synthetic dextrose drop-out medium without Leu and Trp (SD DO-2), then plated on high stringency medium without Leu, Trp, His and Ade (SD DO-4).

### Upregulation of Srs2 protein levels leads to replication stress, chromosome instability and viability loss

We previously observed that *rrp1*+ and *rrp2*+ over-expressing cells accumulated exceptionally bright Rad11 (RPA) foci which may indicate excessive accumulation of single-stranded DNA (ssDNA), as well as fragmented DNA and Rad11 coated bridges [7], which together suggest that these cells suffer chromosome segregation defects. Upregulation of Srs2 also led to the appearance of such foci, but only in about ∼3% of cells (Fig. 4A, B, orange box). The increase in the total Rad11 foci number was also very modest upon Srs2 overexpression, compared to spontaneous foci visible in empty vector control (Fig. 4B). This was accompanied by the accumulation of mitotic aberrations (such as lagging and cut chromosomes), detected by microscopic examination of DAPI-stained cells (Fig. 4C) to a level that was comparable to that reported previously for cells overproducing Rrp1, but much lower than that observed for Rrp2 [7].

**Fig 4.**
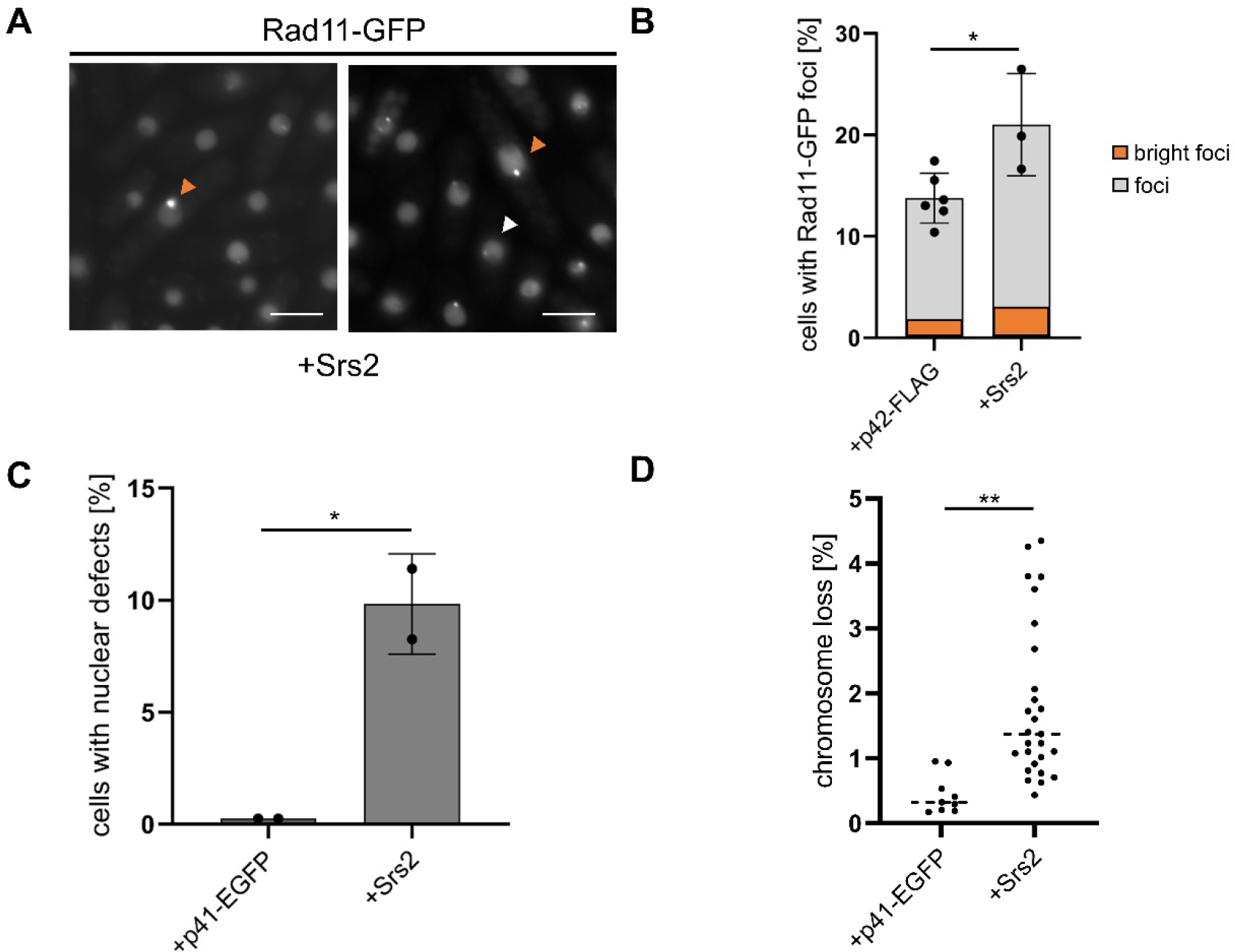
Upregulation of Srs2 protein level leads to chromosome instability. (A) Bright Rad11 foci (examples marked with orange arrowheads) accumulate in *srs2*+ overexpressing cells. The level of regular Rad11 (white arrowheads) also increase (B). The experiment was repeated for at least three independent transformants. Error bars represent the standard deviation about the mean values. Student’s t-test was performed to calculate P-values (* 0.01 < P ≤ 0.05). (C) Mitotic aberrations observed by DAPI staining of the nuclei of transformants grown for 48 h in EMM under expression inducing conditions (-leu) accumulate in cells over-expressing *srs2*+. Two independent transformants for vector and *srs2*+ were analysed and the total number of cells counted was above 900. Error bars represent the deviation from the mean values. Student’s t-test was performed to calculate P-values. (* 0.01 < P ≤ 0.05). (D) Induction of *srs2*+ expression leads to the loss of the nonessential Ch16 minichromosome carrying the ade6-216 allele, resulting in red colony formation on medium with limiting adenine concentration. Error bars represent the deviation from the mean values. Student’s t-test was performed to calculate P-values. (** 0.001 < P ≤ 0.01).

Chromosome instability in cells overproducing Srs2 was assessed using a strain with a nonessential Ch16 mini-chromosome carrying the ade6-216 allele trans-complementing the endogenous ade6-210 allele of the host cell. By scoring the number of red colonies on the medium with a limiting concentration of adenine we observed an increase in the loss of Ch16 mini-chromosome in *srs2*+ over-expressing cells (Fig. 4D), but two fold lower than reported earlier for Rrp1 and Rrp2 overproduction [7].

Overall, the effects of the upregulation of Srs2 are more similar to those for Rrp1 but less severe, especially than those caused by *rrp2*+ over-expression. Together with the fact that overproduced Srs2 seems to localize predominantly in the nucleolus (Fig. 2D), while Rrp1 and Rrp2 are present at centromeres and telomeres [7], the above data indicate that upregulation of Srs2 levels may affects the cells in a way distinct from these translocases.

### Differential checkpoint activation by upregulation of Srs2, Rrp1 and Rrp2

Accumulation of ssDNA and lagging chromosomes that break during mitosis is expected to result in checkpoint activation [21]. We thus reasoned that dysregulation of Srs2, Rrp1, and Rrp2 protein levels should result in DNA damage and/or replication checkpoint pathway activation. This can be readily observed in *S. pombe* as an increase in cell length and indeed cells over-expressing *srs2*+ as well as *rrp1*+ or *rrp2*+ were elongated (Fig. 5A). Rad3, a protein related to mammalian ATR, is a key *S. pombe* kinase activated upon DNA damage or replication arrest, and phosphorylates its effector kinases, Chk1 and Cds1, responsible for activation of DNA damage and replication checkpoints, respectively [22]. Rad3 is also responsible for phosphorylation of histone H2A in fission yeast in response to DNA damage [23] but also when replication fork stall at various barriers, repetitive DNA and heterochromatin in the centromeres and telomeres, as well as ribosomal DNA (rDNA) [24]. We found that in cells over-expressing *srs2*+ as well as *rrp1*+ or *rrp2*+ the level of γH2A was increased (Fig. 5B), thus confirming the activation of the checkpoint response.

**Fig. 5.**
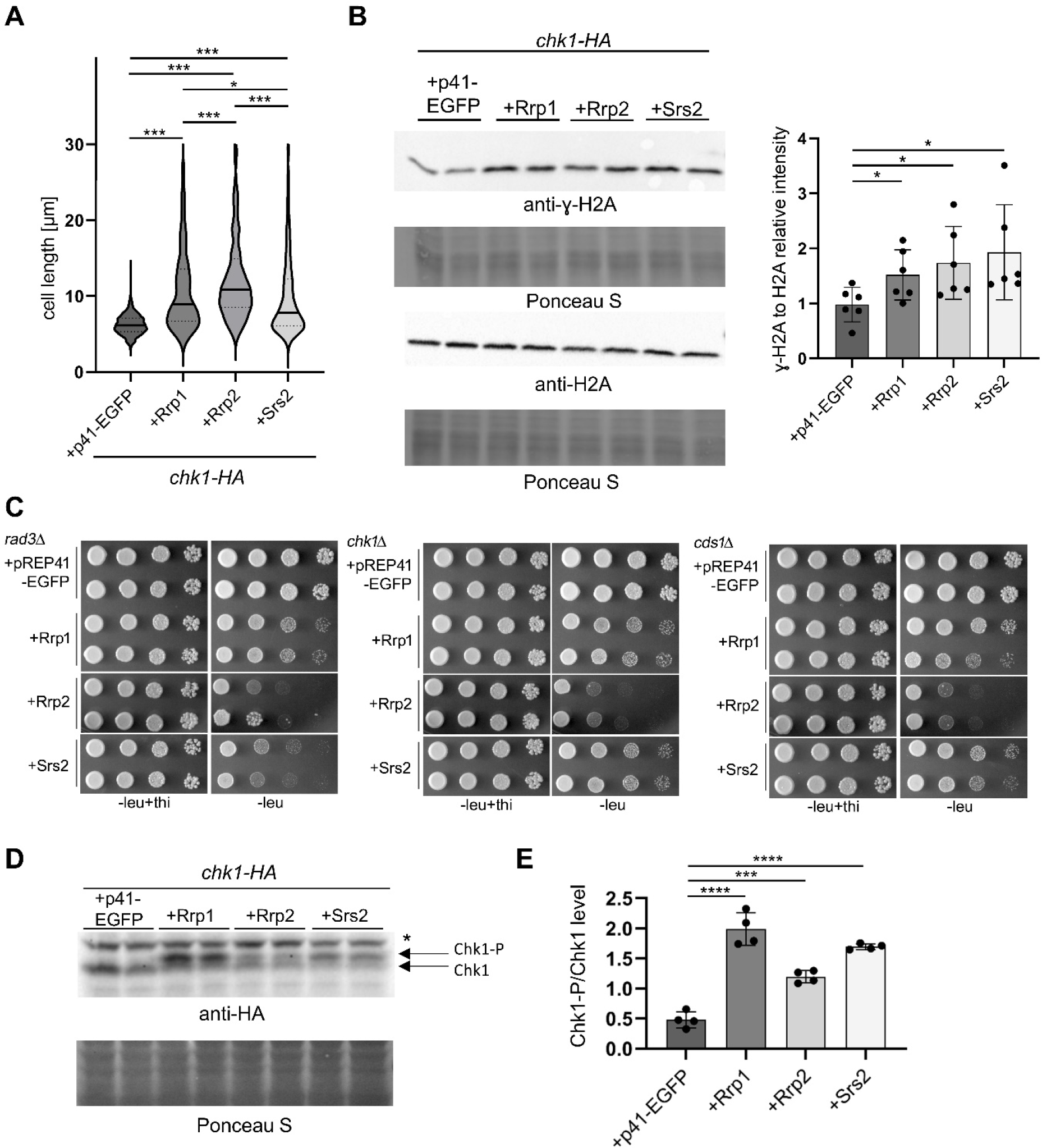
Checkpoint response to upregulation of Srs2, Rrp1 and Rrp2. (A) Over-expression *srs2*+, *rrp1*+, *rrp2*+ leads to the increase in cell length. Respective transformants in Chk1-HA strain were grown for 48h in EMM under expression inducing conditions (-leu), observed under the microscope and analysed with Axiovision rel. 4.8. Data from two independent transformations were analysed and the total number of cells counted was above 375.The centre line represents the median; the dotted lines represent upper and lower quartiles. Student’s t-test was performed to calculate P-values (*** P ≤ 0.001, * 0.01 < P ≤ 0.05). (B) γH2A phosphorylation increases upon overproduction of all studied proteins. Total protein extracts were collected after 24h growth in EMM-leu to induce *nmt* promoter and analysed by Western blot using anti-γH2A and anti-H2A antibodies. Data were quantified and shown as the intensity of γH2A signal versus H2A signal. Reads were normalised by the mean value obtained for vector control samples. Western blots from at least two separate protein isolations from two different transformants were examined. Error bars represent the standard deviation about the mean values. Student’s t-test was performed to calculate P-values (* 0.01 < P ≤ 0.05). (C) Deletion of *rad3+* only aggravated the growth defect induced by *srs2*+ over-expression. Cells of respective transformants were appropriately diluted and spotted on EMM plates where expression of respective genes was induced (-leu) or repressed (-leu + thi), incubated for 6 days and photographed. (D) Over-expression *srs2*+, *rrp1*+, *rrp2*+ leads to the increase in Chk1 phosphorylation. Total protein extracts were collected after 24h induction of *nmt* promoter and analysed by Western blot. Chk1 was detected with anti-HA antibody and its phosphorylated form is visible as slower migrating band (Chk1-PO4, unspecific band is marked as *). (E) Data were quantified and shown as intensity of Chk1-PO4 signal, versus Chk1 signal. Reads were normalised by the mean value obtained for vector control samples. The experiment was repeated at twice for two independent transformants. Error bars represent the standard deviation about the mean values. Student’s t-test was performed to calculate P-values (**** P ≤ 0.0001, *** P ≤ 0.001).

Deletion of *rad3+* aggravated the growth defect induced by *srs2*+ over-expression, however deletion of *cds1*+ or *chk1*+ had no visible effect (Fig. 5C), indicating that DNA damage and replication checkpoint pathways may have redundant roles in augmenting the survival of cells overproducing Srs2. This is different to what has been observed in *S. cerevisiae* where toxicity of *SRS2* over-expression significantly increased when DNA replication, but not DNA damage, checkpoint was compromised [19]. Growth defect induced by Rrp1 and Rrp2 overproduction in all 3 checkpoint mutants was comparable to that in wild-type cells (Fig.5C). This indicates that the checkpoint response might be affected in different ways upon upregulation of Srs2, Rrp1 and Rrp2.

Indeed, we observed that *srs2*+, *rrp1*+ or *rrp2*+ over-expression resulted in Chk1 phosphorylation, albeit to different levels (Fig. 5D) that did not correlate with severity of growth defect associated with increased levels of these proteins seen in Fig. 3A, suggesting different contributions of DNA damage and replication stress response to checkpoint activation. The toxicity of *rrp2+* overexpression was the most severe, yet resulted in the mildest activation of Chk1. In contrast, phosphorylation of Chk1 was most apparent in *rrp1+* over-expressing cells, with Srs2 overproduction having the intermediate effect (Fig. 5E).

We thus decided to assess the relative contribution of Chk1 and Cds1 pathways by measuring the cell length of respective checkpoint mutants, overproducing Srs2, Rrp1 or Rrp2 proteins as compared to wild-type. As expected, we found that in *rad3*Δ mutant over-expression of all three proteins failed to induce checkpoint activation measured as an increase in cell length relative to empty vector control (Fig. 6A-C). Cell length was decreased to a similar degree in *chk1*Δ and *cds1*Δ mutants over-expressing *srs2*+ (Fig. 6A) suggesting that overproduction of Srs2 activated both DNA damage and replication stress response pathways. However, checkpoint activation seemed to depend mostly on DNA damage effector kinase Chk1 upon *rrp1*+ over-expression (Fig. 6B). Checkpoint response for *rrp2*+ over-expressing cells was more dependent on the replication stress effector kinase Cds1 (Fig. 6C). These results are consistent with levels of Chk1 phosphorylation observed: intermediate when Srs2 was overproduced in wild-type cells, highest for Rrp1 and lowest for Rrp2 overproduction (Fig. 5D-E).

**Fig 6.**
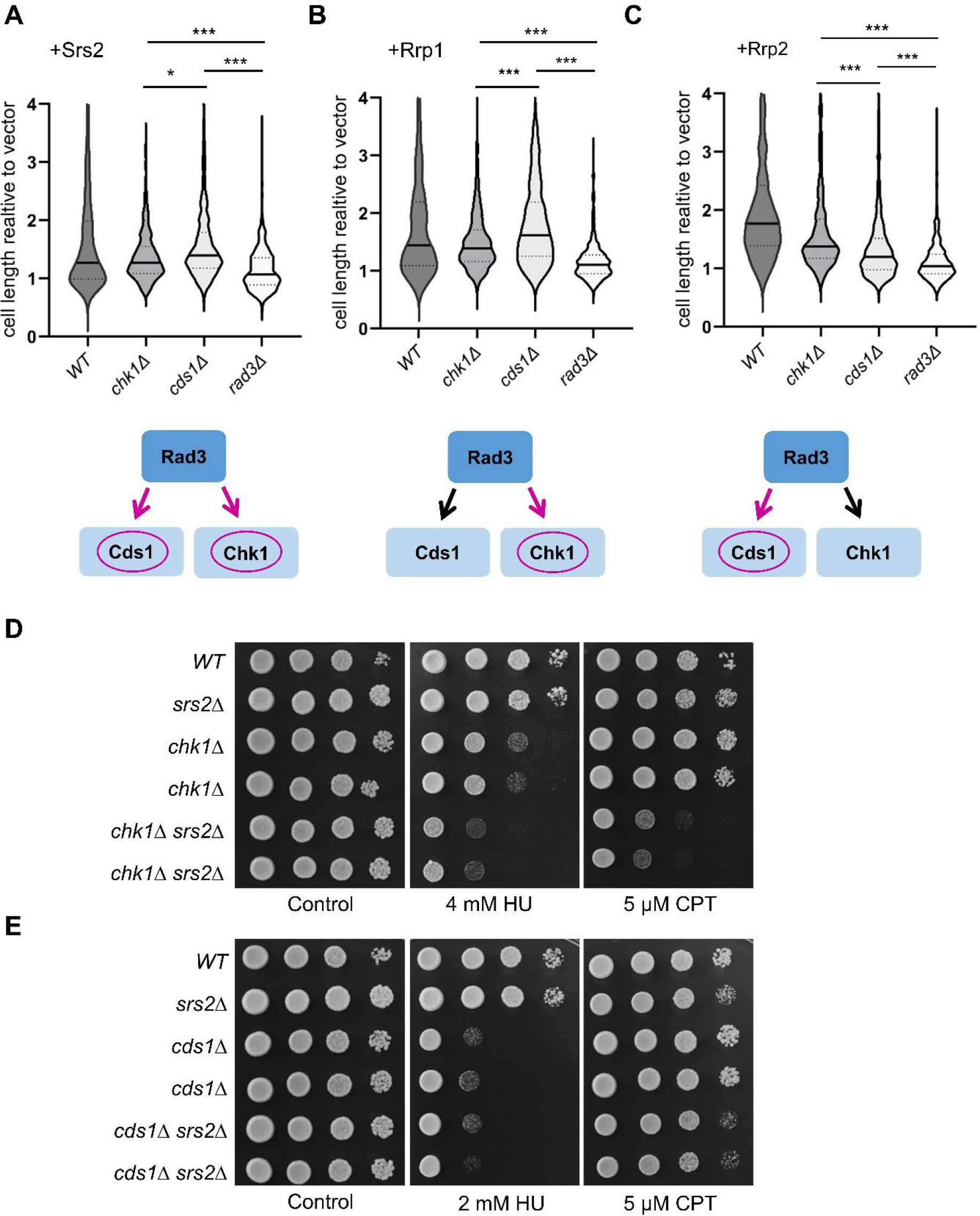
DNA damage and replication stress checkpoint pathways are differently activated by upregulation of Srs2, Rrp1 and Rrp2 protein levels. Lack of Rad3, Chk1 and Cds1 differentially affects the length of cells over-expressing *srs2*+ (A), *rrp1*+ (B) and *rrp2*+ (C). Diagrams depicting major activated pathway are shown below. Transformants in respective deletion were grown for 48h in EMM under expression inducing conditions (-leu), observed under the microscope and analysed with Axiovision rel. 4.8. Data from two independent transformations were analysed and the total number of cells counted was above 375. The centre line represents the median; the dotted lines represent upper and lower quartiles. Student’s t-test was performed to calculate P-values (*** P ≤ 0.001, * 0.01 < P ≤ 0.05). Epistasis between srs2+ helicase and *chk1*+ (D) and *cds1*+ (E) genes. Cultures of respective single and *chk1*Δ*srs2*Δ and *cds1*Δ*srs2*Δ double mutant strains were appropriately diluted, spotted on YES plates with HU or CPT, incubated for 4 days and photographed.

Srs2 has been shown to work in a pathway redundant to Rad3 and Mrc1, possibly independent from checkpoint signalling functions of these proteins but requiring Srs2 ATPase activity [20]. Consistently, we observed that the double *chk1*Δ*srs2*Δ mutant was more sensitive to HU and CPT than single *chk1*Δ mutant (Fig. 6D), while no additive effect on sensitivity for double *cds1*Δ*srs2*Δ mutant was seen (Fig. 6E). This suggests that Srs2 has an important role in preventing DNA damage during replication and this becomes especially apparent when DNA damage checkpoint pathway cannot be activated.

## Discussion

Rrp1 and Rrp2 have been shown to be required for the full rescue of the *rqh1*Δ mutant’s HU sensitivity and aberrant mitosis phenotype by deletion of *rad57+* and proposed to act together with Swi5 and Srs2 in a synthesis-dependent strand annealing HR repair pathway [5]. Here we show that this rescue is also dependent on the presence of Srs2 supporting our published model placing some aspects of Srs2 activity in a common pathway with the Rrp1-Rrp2 complex.

Subsequently, by analysing the effect of upregulation of Rrp1 and Rrp2 protein levels we have found that they have activities independent from one another. Rrp1 is more important for centromere maintenance and its translocase and ubiquitin ligase activities are crucial for the regulation of Rad51 recombinase [7,9], while Rrp2 has been shown to protect cells from Top2-induced DNA damage and be required for telomere stability [7,8]. Here we set out to analyse the effect of Srs2 overproduction.

We found that upregulation of Srs2 protein levels is associated with mild accumulation of ssDNA seen as an increase in the number of bright Rad11-EGFP foci, anaphase aberrations, chromosome instability and viability loss. These effects are similar to those seen for upregulation of Rrp1 levels but less severe than those caused by *rrp2*+ overexpression. However, Srs2 localisation in the nucleus and checkpoint response to its overproduction appears to be different from Rrp1 and Rrp2.

We have previously demonstrated that overproduced Rrp1 and Rrp2 localise as distinct foci within the nucleus, and have been shown to be most abundant at centromeres and telomeres, respectively. Consistently, their activity was especially important during replication stress response at these respective regions [7]. Our yeast-two-hybrid experiments suggest that Srs2 does not form a direct complex with either Rrp1 or Rrp2. Moreover, fluorescence microscopy analyses presented here suggest that Srs2 mainly localises to the rDNA region within nucleus. In line with this localization Srs2 has been previously shown to be crucial for rDNA maintenance redundantly with the non-checkpoint function of Mrc1, a mediator of the replication checkpoint pathway [20]. Moreover, enhancing fork stalling at replication barriers has been demonstrated to be a replication checkpoint independent function of Mrc1 [25]. Interestingly, Srs2 has been proposed to be required for a slow restart of DNA replication arrested at *RTS1*, a polar barrier present at *S. pombe* mating-type locus, but related to barriers found in the rDNA gene arrays as well as centromeres and telomeres [26]. We have shown that cells lacking Srs2 rely on functional DNA damage checkpoint for replication stress response as the double *chk1*Δ*srs2*Δ mutant was more sensitive to HU and CPT than the single *chk1*Δ mutant. One may thus hypothesize that Srs2 may participate in regulating the timing and speed of restart of replication forks stalled at repeats within the rDNA region, in order to prevent unscheduled events leading to DNA damage and the activation of Chk1-dependent checkpoint pathway. Upon upregulation of Srs2 levels this process would be disturbed.

It would be interesting in the future to determine to what extent the observed differences in checkpoint activation, as well as similarities and dissimilarities in other phenotypes induced by overproduction of Srs2, Rrp1 and Rrp2 described above, result from the differences in the place of action of these three proteins as well as their interacting partners. This would shed light on the specific requirements for the regulation of replication stress response pathways in different difficult to replicate regions in *S. pombe* genome.

## Acknowledgments

We thank Matthew C. Whitby, Tony M. Carr, Jo Murray and Hiroshi Iwasaki for providing strains and to Ireneusz Litwin for comments on the manuscript.

## Author contributions

Gabriela Baranowska, Investigation, Formal analysis, Funding acquisition, Writing - review and editing; Dorota Misiorna Investigation, Formal analysis, Methodology, Writing - review and editing; Wojciech Białek, Investigation, Formal analysis, Methodology; Karol Kramarz, Investigation, Formal analysis, Methodology, Supervision, Funding acquisition, Writing - original draft, Writing - review and editing; Dorota Dziadkowiec, Conceptualization, Supervision, Formal analysis, Writing - original draft, Writing - review and editing.

## Supporting information

**S1 Table. Strains used in this study**.

**S2 Table. Plasmids used in this study**.

**S3 Table. Primers used in this study**.

