## Supplementary Tables 1-3 for "Replication stress response in fission yeast differentially depends on maintaining proper levels of Srs2 helicase and Rrp1, Rrp2 DNA translocases"

**Supplementary files** Baranowska G., *et al.* 2023

**S1 Table. Strains used in this study.**

| Strain | Genotype | Source |
|---|---|---|
| YA254 (WT) | ura4-D18, leu1-32, his3-D1, arg3-D1, h90 | a |
| *rrp1*Δ | rrp1D:kanMX6, ura4-D18, leu1-32, his3-D1, arg3-D1, h90 | b |
| *rrp2*Δ | rrp2D::kanMX6, ura4-D18, leu1-32, his3-D1, arg3-D1, h90 | b |
| *rrp1*Δ | rrp1D::natMX6, his3-D1, leu1-32, ura4-D18, arg3-D1, h90 | b |
| *rrp2*Δ | rrp2D::natMX6, his3-D1, leu1-32, ura4-D18, arg3-D1, h90 | b |
| *srs2*Δ | srs2D::ura4+, ura4-D18, his3-D1, leu1-32, arg3-D4 h90 | b |
| *rqh1*Δ | rqh1D::ura4+, ura4-D18, his3-D1, leu1-32, arg3-D4 h90 | b |
| *rad57*Δ | rad57D::his3+, ura4-D18, leu1-32, his3-D1, arg3-D1, smt0 | a |
| *rqh1*Δ*rhp57*Δ | rqh1D::ura4+, rhp57D::his3+, ura4-D18, his3-D1, leu1-32, arg3-D4 smt0 | b |
| *rqh1*Δ*rhp57*Δ*rrp1*Δ | rqh1D::ura4+, rhp57D::his3+, rrp1D:*kan*MX6, ura4-D18, his3-D1, leu1-32, arg3-D4 smt0 | b |
| *rqh1*Δ*rhp57*Δ*rrp2*Δ | rqh1D::ura4+, rhp57D::his3+, rrp2D:*kan*MX6, ura4-D18, his3-D1, leu1-32, arg3-D4, smt0 | b |
| *rqh1*Δ*rhp57*Δ*srs2*Δ | srs2D::ura4+, rhp57::his3+, rqh1D:*kan*MX6 ura4-D18, his3-D1, leu1-32, arg3-D4, smt0 | c |
| *rrp1*Δ*srs2*Δ | srs2D::ura4+, rrp1D::*kan*MX6, leu1-32, ura4-D18, his3-D1, h90 | b |
| *rrp2*Δ*srs2*Δ | srs2D::ura4+, rrp2D::*kan*MX6, leu1-32, ura4-D18, his3-D1, arg3-D4, h90 | b |
| *rad57*Δ | rad57D::his3+, ura4-D18, leu1-32, his3-D1, arg3-D1, Msmt0 | a |
| *rad51*Δ | rhp51D::his3+, ura4-D18, his3-D1, leu1-32, arg3-D4 smt0 | a |
| *rad3*Δ | rad3D::kanMX6, ura4-D18, leu1-32, ade6-704, h- | d |
| *cds1*Δ | cds1D::natMX6, ura4-D18, leu1-32, ade6-704, h- | d |
| *chk1*Δ | chk1D::kanMX6, ura4-D18, leu1-32, ade6-704, h- | d |
| *cds1*Δ*srs2*Δ | cds1D::natMX6, srs2D::ura4+, ura4-D18, leu1-32, ade6-704, his3-D1, arg3-D1, h- | c |
| *chk1*Δ*srs2*Δ | chk1D::kanMX6, srs2D::ura4+, ura4-D18, leu1-32, ade6-704, his3-D1, arg3-D1, h- | c |
| AW161 | chk1-HA, ura4-D18, leu1-32, his3-D1, h+ | d |
| CJ01 | ura4-D18; leu1-32; his3-D1; arg3-D1; ade6-m210; Chr16 ade6-m216; h+ | e |
| AMC377 | rad11-GFP::kanMX6, ura4-D18, leu1-32, his3-D1, h+ | d |
| AH 109 | MATa, trp1-901, leu2-3, 112, ura3-52, his3-200, gal4D, gal80D, LYS2::$GAL1_{UAS}$-$GAL1_{TATA}$-HIS3, $GAL2_{UAS}$-$GAL2_{TATA}$-ADE2, URA3::$MEL1_{UAS}$- $MEL1_{TATA}$-lacZ | Clontech |

a - Hiroshi Iwasaki
b - laboratory stock
c - this work
d - Tony M Carr
e - Jo Murray

**S2 Table. Plasmids used in this study.**

| Plasmid | Source |
|---|---|
| pREP81-FLAG | a |
| pREP81-Rrp1-FLAG | a |
| pREP81-Rrp2-FLAG | a |
| pREP81-Srs2-FLAG | b |
| pREP41-EGFP | c |
| pREP41-EGFP-Rrp1-FLAG | b |
| pREP41-EGFP-Rrp2-FLAG | b |
| pREP41-EGFP-Srs2-FLAG | a |
| pREP42-HA | c |
| pREP42-HA-Rrp1 | a |
| pREP42-HA-Rrp2 | a |
| pREP42-HA-Srs2-FLAG | b |

a - laboratory stock
b - this study
c - Craven,R.A., *et al.* (1998) Gene, 221, 59–68. doi:10.1016/s0378-1119(98)00434-x

**S3 Table. Primers used in this study.**

| Cloned gene | Primer name | Primer sequence |
|---|---|---|
| FLAG | REP3flag_a_r<br>REP3flag_b_f<br>REP3flag_c_r<br>REP3flag_d_f | ctttatcatcgtcgtccttgtagtcggatcctctagagtcgacatatgatttaac<br>tacaaggacgacgatgataaagactacaaggacgacgatgataaagacta<br>gggtcatttatcatcgtcgtccttgtagtctttatcatcgtcgtccttgtagt<br>caaggacgacgatgataaatgacccgggtaaaaggaatgtctcccttgccagtac |
| Srs2-FLAG | srsEx1_fwd<br>srsEx1_rev<br>srsEx2_fwd<br>srsEx2_rev<br>srsEx3_fwd<br>srsEx3_rev<br>srsEx4_fwd<br>srsEx4_rev | atagtcgctttgttaaatcatATGGAAACGAAATCATCATAC<br>aatagcttgaGTATCATTTCTGTCAGCAATC<br>gaaatgatacTCAAGCTATTATGAAGCGAC<br>catcaaaatcCGCTAAATTGTTCTTCCATAAG<br>caatttagcgGATTTTGATGATTTGCTTTTAAACTTTATTTTATTAC<br>gtttaatttctCTTGCGATCCAGTAAGATTC<br>ggatcgcaagAGAAATTAAACGTATAGTAGGTTC<br>catcgtcgtccttgtagtcggatccTAACATTCGTGAAACTCGTAG |
